# Deep Recurrent Neural Network Reveals a Hierarchy of Process Memory during Dynamic Natural Vision

**DOI:** 10.1101/177196

**Authors:** Junxing Shi, Haiguang Wen, Yizhen Zhang, Kuan Han, Zhongming Liu

## Abstract

The human visual cortex extracts both spatial and temporal visual features to support perception and guide behavior. Deep convolutional neural networks (CNNs) provide a computational framework to model cortical representation and organization for spatial visual processing, but unable to explain how the brain processes temporal information. To overcome this limitation, we extended a CNN by adding recurrent connections to different layers of the CNN to allow spatial representations to be remembered and accumulated over time. The extended model, or the recurrent neural network (RNN), embodied a hierarchical and distributed model of process memory as an integral part of visual processing. Unlike the CNN, the RNN learned spatiotemporal features from videos to enable action recognition. The RNN better predicted cortical responses to natural movie stimuli than the CNN, at all visual areas especially those along the dorsal stream. As a fully-observable model of visual processing, the RNN also revealed a cortical hierarchy of temporal receptive window, dynamics of process memory, and spatiotemporal representations. These results support the hypothesis of process memory, and demonstrate the potential of using the RNN for in-depth computational understanding of dynamic natural vision.

## INTRODUCTION

Human behavior depends heavily on vision. The brain’s visual system works efficiently and flexibly to support a variety of tasks, such as visual recognition, tracking, and attention, to name a few. Although a computational model of natural vision remains incomplete, it has evolved from shallow to deep models to better explain brain activity (Kriegeskorte, 2015; Khaligh-Razavi et al., 2017), predict human behaviors (Canziani and Culurciello, 2015; Fragkiadaki et al., 2015; Mnih et al., 2015), and support artificial intelligence (AI) (LeCun et al., 2015; Silver et al., 2016). In particular, convolutional neural networks (CNNs), trained with millions of labeled natural images (Russakovsky et al., 2015), have enabled computers to recognize images with human-like performance (He et al., 2015). CNNs bear similar representational structures as the visual cortex (Khaligh-Razavi and Kriegeskorte, 2014; Cichy et al., 2016) and predict brain responses to natural stimuli (Yamins et al., 2014; Güçlü and van Gerven, 2015a; Wen et al., 2016, 2017a; Eickenberg et al., 2017). It thus provides new opportunities for understanding cortical representations of vision (Yamins and DiCarlo, 2016; Khaligh-Razavi et al., 2017).

Nevertheless, CNNs driven for image recognition are incomplete models of the visual system. CNNs are intended and trained for analyses of images in isolation, rather than videos where temporal relationships among individual frames carry information about action. In natural viewing conditions, the brain integrates information not only in space (Hubel and Wiesel, 1968) but also in time (Hasson et al., 2008). Both spatial and temporal information is processed by cascaded areas with increasing spatial receptive fields (Wandell et al., 2007) and temporal receptive windows (TRWs) (Hasson et al., 2008) along the visual hierarchy. It has been hypothesized that temporal processing requires cortical circuits to maintain distributed “process memory”, which allows information to be accumulated over a variety of timescales (Hasson et al., 2015). However, CNNs only model spatial processing via feedforward-only computation, lacking any mechanism for processing temporal information.

An effective way to model temporal processing is by using recurrent neural networks (RNNs), which learn representations from sequential data (Goodfellow et al., 2016). As its name indicates, a RNN processes the incoming input by also considering its own output given the history input. In AI, RNNs have made impressive progress in speech and action recognition (Jozefowicz et al., 2015; Greff et al., 2016; Donahue et al., 2015), demonstrating the potential to match the human performance on such tasks. In addition, RNN can be designed with an architecture that resembles the notion of “process memory”, as recently proposed in neuroscience (Hasson et al., 2015). Therefore, RNN is a logical step forward from CNN toward modeling and understanding the inner working of the visual system in dynamic natural vision.

In this study, we designed, trained, and tested a RNN to model and explain cortical processes for spatial and temporal visual processing. This model began with a static CNN pre-trained for image recognition (Simonyan and Zisserman, 2014a). Recurrent connections were added to different layers in the CNN to embed process memory into spatial processing, so that layer-wise spatial representations could be remembered and accumulated over time to form video representations. While keeping the CNN intact and fixed, the parameters for the recurrent connections were optimized by training the entire model for action recognition with a large set of labeled videos (Soomro et al., 2012). Then, we evaluated how well this RNN model matched the human visual cortex up to linear transform. Specifically, the RNN was trained to predict functional magnetic resonance imaging (fMRI) responses to natural movie stimuli. The prediction accuracy with the RNN was compared with that of the CNN, to address whether and where the recurrent connections allowed the RNN to better model cortical representations given dynamic natural stimuli. Through the RNN, we also characterized and mapped the cortical topography of temporal receptive windows and dynamics of process memory. By doing so, we attempted to use a fully-observable model of process memory to explain the hierarchy of temporal processing, as a way to directly test the hypothesis of process memory (Hasson et al., 2015).

## METHODS AND MATERIALS

### Experimental Data

The experimental data was from our previous studies (Wen et al., 2016, 2017a, b), according to a research protocol approved by the Institutional Review Board at Purdue University. Briefly, we acquired fMRI scans from three healthy subjects while they were watching natural videos. The video-fMRI data was split into two datasets to train and test the encoding models, respectively, for predicting fMRI responses given any natural visual stimuli. The training movie contained 12.8 hours of videos for Subject 1, and 2.4 hours for the other subjects (Subject 2&3). The testing movie for every subject contained 40 minutes of videos, presented ten times during fMRI. These movies included a total of ∼9,300 continuous videos without abrupt scene transitions, covering a wide range of realistic visual experiences. These videos were concatenated and then split into 8-min movie sessions, each of which was used as the stimuli in a single fMRI experiment. Subjects watched each movie session through a binocular goggle (20.3^o^×20.3^o^) with their eyes fixating at the center of the screen (red cross). Whole-brain fMRI scans were acquired in 3-T with an isotropic resolution of 3.5mm and a repetition time of 2s. The fMRI data was preprocessed and co-registered onto a standard cortical surface (Glasser et al., 2013). More details about the stimuli, data acquisition, and preprocessing are described in (Wen et al., 2016, 2017a).

### Convolutional Neural Network (CNN)

Similar to our prior studies (Wen et al., 2016, 2017a, b), a pre-trained CNN, also known as the VGG16 (Simonyan and Zisserman, 2014a), was used to extract the hierarchical feature representations of every video frame as the outputs of artificial neurons (or units). This CNN contained 16 layers of units stacked in a feedforward network for processing the spatial information in the input. Among the 16 layers, the first 13 layers were divided into five blocks (or sub-models). Each block started with multiple convolutional layers with Rectified Linear Units (ReLU) (Nair and Hinton, 2010), and ended with spatial max-pooling (Boureau et al., 2010). To simplify terminology, hereafter we refer to these blocks as layers. The outputs from every layer were organized as three-dimensional arrays (known as feature maps). For the 1^st^ through 5^th^ layers, the sizes of feature maps were 64×112×112, 128×56×56, 256×28×28, 512×14×14, and 512×7×7, where the 1^st^ dimension was the number of features, the 2^nd^ and 3^rd^ dimensions specified the width and the height (or the spatial dimension). From lower to higher layers, the number of features increased as the dimension per feature decreased. This CNN was implemented in *PyTorch* (http://pytorch.org/).

### Recurrent Neural Network (RNN)

A RNN was constructed by adding recurrent connections to the four out of five layers in the CNN. The first layer was excluded to reduce the computational demand as in a prior study (Ballas et al., 2015). The recurrent connections served to model distributed process memory (Hasson et al., 2015), which allowed to memorize and accumulate visual information over time for temporal processing.

**Fig. 1A** illustrates the design of the RNN for extracting layer-wise feature representations of an input video. Let the input video be a time series of color (RGB) frames with 224×224 pixels per frame. For the video frame **x**_*t*_ at time *t*, **x**_*t*_ ∈ ℝ^3×224×224^. The internal states of the RNN at layer *l*, denoted as 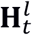, was updated at each moment, according to the incoming information **x**_*t*_ and the history states 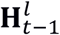, as expressed in Eq. (1).

**Figure 1.**
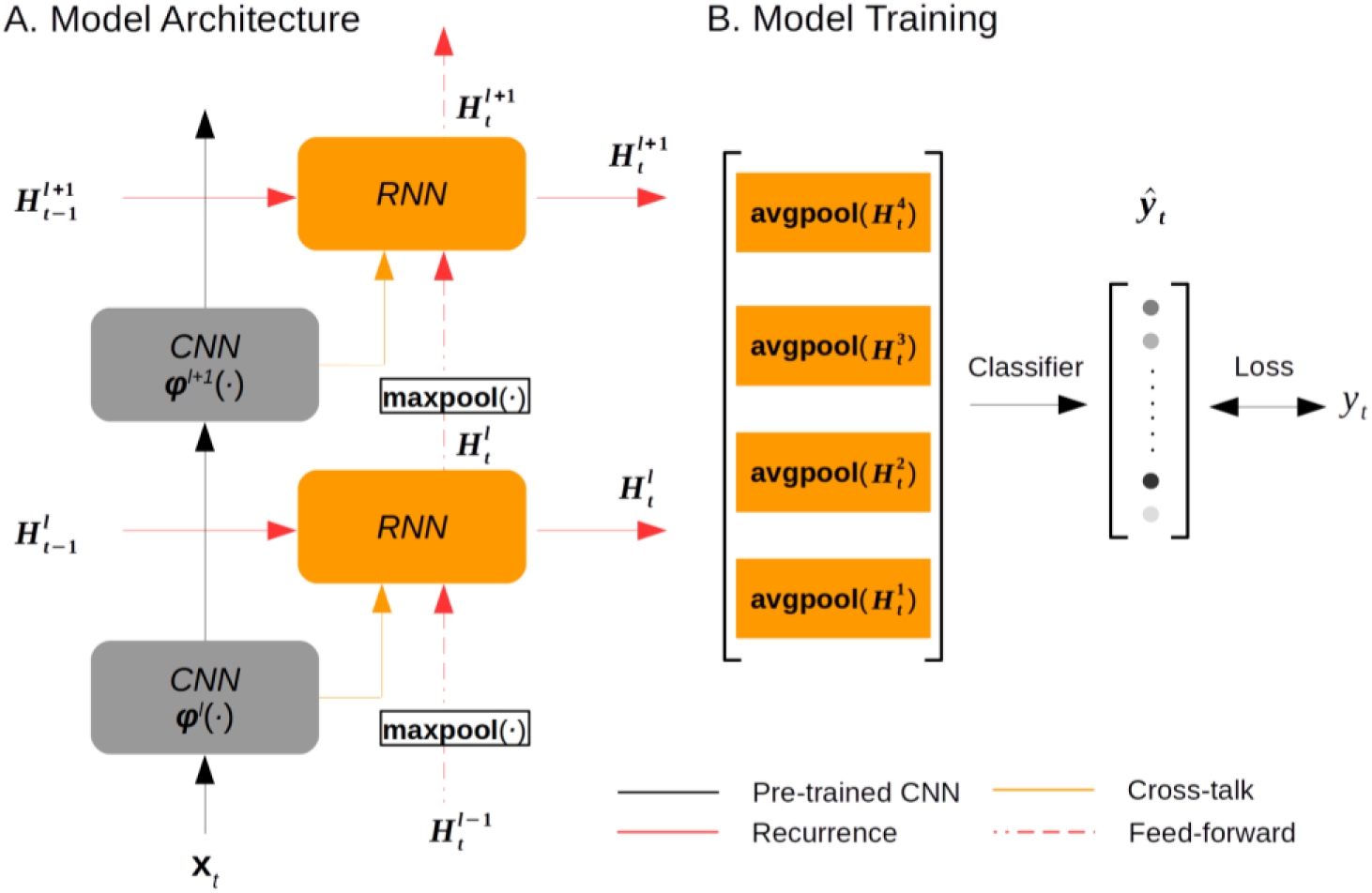
The recurrent model of vision. **A)** The architectural design of the RNN. **B**) The model training strategy. The gray blocks indicate the CNN layers; the orange blocks indicate the RNN layers. The CNN was pre-trained and fixed, while the RNN was optimized on the task of action recognition.

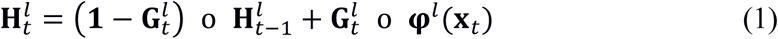

where **ψ**^*l*^(·) was the spatial features encoded at layer *l* in the pre-trained CNN, so **ψ**^*l*^(**x**_*t*_) was the extracted feature representations of the current input **x**_*t*_. Importantly, 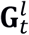 was the so-called “forget gate” essential to learning long-term temporal dependency (Pascanu et al., 2013). As its name indicates, the forget gate determined the extent to which the history states should be “forgotten”, or reversely the extent to which the incoming information should be “remembered”. As such, the forget gate controlled, moment by moment, how information should be stored into vs. retrieved from process memory. Given a higher value of the forget gate, the RNN’s current states 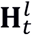 were updated by retrieving less from its “memory” 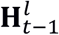, but learning more from the representations of the current input **ψ**^*l*^(**x**_*t*_). This notion was expressed as the weighted sum of the two terms in the right-hand side of Eq. (1), where o stands for Hadamard product and the weights of the two terms sum to 1. In short, the RNN embedded an explicit model of the “process memory” (Hasson et al., 2015).

Note that the forget gate 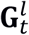 was time dependent but a function of the time-invariant weights, denoted as **ω**^*l*^, of the recurrent connections, expressed as below.

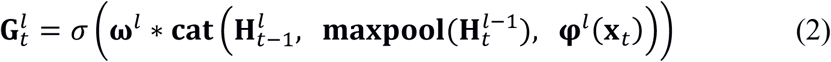

where σ(·) is the sigmoid function whose output ranges from 0 to 1.

As expressed in Eq. (2), the forget gate 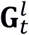 was the weighted sum of three terms: the RNN’s previous output 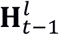, the CNN’s current output **ψ**^*l*^(**x**_*t*_), and the RNN’s current input from the lower layer **maxpool**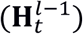. Here, **maxpool**(•) stands for the max-pooling operation, which in this study used a kernel size of 2×2 and a stride of 2 to spatially subsample half of the RNN’s output at layer *l-1* and fed the result as the input to layer *l* in the RNN. Note that the weighted summation was in practice implemented as convolving a 3×3 kernel (with a padding of 1) across all three input terms concatenated together, as expressed by **cat**(•) in Eq. (2). This reduced the number of unknown parameters to be trained. In other words, **ω**^*l*^ ∈ ℝ^M×N×3×3^, where M and N were the numbers of output and input feature maps, respectively.

### Training the RNN for Action Recognition

The RNN was trained for video action recognition by using the first split of the UCF101 dataset (Soomro et al., 2012). The dataset included 9,537 training videos and 3,783 validation videos from 101 labeled action categories. All videos were resampled at 5 frames per second, and preprocessed as described elsewhere (Ballas et al., 2015), except that we did not artificially augment the dataset with random crops.

To train the RNN with labeled action videos, a linear softmax classifier was added to the RNN to classify every training video frame as one of the 101 action categories. As expressed by Eq. (3), the inputs to the classifier were the feature representations from all layers in the RNN, and its outputs were the normalized probabilities, by which a given video frame was classified into pre-defined categories (**Fig. 1B**).

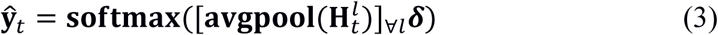

where **avgpool**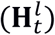 reduced 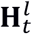 from a 3-D feature array to a 1-D feature vector by averaging over the spatial dimension (or average pooling); [·]_∀***l***_ further concatenated the feature vectors across all layers in the RNN; ***δ*** ∈ **R**^Px101^ was a trainable linear function to transform the concatenated feature vector onto a score for each category; **softmax**(·) converted the scores into a probability distribution, 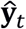, to report the result of action categorization given each input video frame.

The loss function for training the RNN was defined as below.

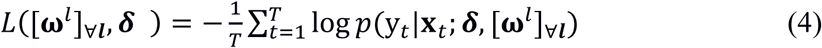

where y_*t*_ stands for the true action category labeled for the input **x**_*t*_. Here, the learning objective was to maximize the average (over *T* samples) log probability of correct classification conditioned on the input {**x**_*t*_}_∀*t*_ and parameterized by linear projection ***δ*** and the recurrent parameters [**ω**^*l*^]_∀*l*_.

The RNN was trained by using mini-batch gradient descent and back-propagation through time (Werbos, 1990). The parameters [**ω**^*l*^]_∀***l***_ were initialized as random values from a uniform distribution between –0.01 and 0.01. For the training configurations, the batch size was set to 10. The sequence length was 20 frames, so that the losses were accumulated over 20 consecutive frames before back-propagation. A dropout of 0.7 was used to train ***δ***. The gradient vector was normalized to 5. The gradient descent algorithm was based on the Adam optimizer (Kingma and Ba, 2014) with the learning rate initialized as 1e-3. The learning rate was decayed by 0.1 every 10 epochs, while the learning iterated across all training videos in each epoch.

To evaluate the RNN on the task of action recognition, we evaluated the top-1 accuracy given the validation videos, while being top-1 accurate meant that the most probable classification matched the label. In addition, we also trained a linear softmax classifier based on the feature representations extracted from the CNN with the same training data and learning objective, and evaluated the top-1 accuracy for model comparison.

### Encoding Models

For each subject, a voxel-wise encoding model (Naselaris et al., 2011) was established for predicting the fMRI response to natural movie stimuli based on the features of the movie extracted by the RNN (or the CNN for comparison). A linear regression model was trained separately for each voxel to project feature representations to voxel responses, similar to prior studies (Güçlü and van Gerven, 2015a, b; Wen et al., 2016, 2017a, b; Eickenberg et al., 2017). As described below, the same training methods were used regardless of whether the RNN or the CNN was used as the feature model.

Using the RNN (or the CNN), the feature representations of the training movie were extracted and sampled every second. Note that the feature dimension was identical for the CNN and the RNN, both including feature representations from four layers with exactly matched numbers of units in each layer. For each of the four layers, the number of units was 401408, 200704, 100352, and 25088. Combining features across these layers ended up with a very high-dimensional feature space. To reduce the dimension of the feature space, principal component analysis (PCA) was applied first to each layer and then to all layers, similar to our prior studies (Wen et al., 2016, 2017a, 2017b). The principal components (PCs) were identified based on the feature representations of the training movie, and explained 90% variance. Such PCs defined a set of orthogonal basis vectors, spanning a subspace of the original feature space (or the reduced feature space). Applying this basis set as a linear operator, **B**, to any representation, **X**, in the original feature space, converted it to the reduced feature space, as expressed as Eq. (5).; applying the transpose of **B** to any representation, **Z**, in the reduced feature space, converted it to the original feature space.

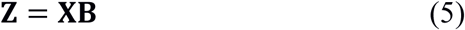

where **X** ∈ ℝ^*T*×*q*^ stands for the representation of the RNN (or the CNN) with *T* samples and *q* units; **B** is a *q*-by-*p* matrix that consists of the PCs identified with the training movie; and **Z** ∈ ℝ^*T*×*p*^ stands for the *p*-dimensional feature representations after dimension reduction (*p<q*).

The feature representations after dimension reduction (i.e. columns in **Z**) were individually convolved with a canonical hemodynamic response function (HRF) (Buxton et al., 2004) and downsampled to match the sampling rate for fMRI. Then, **Z** was used to fit each voxel’s response during the training movie through a voxel-specific linear regression model, expressed as Eq. (6).

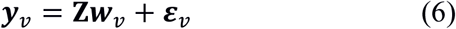

where ***w***_*v*_ is a columnar vector of regression coefficients specific to voxel *v*, and ε_*v*_ is the error term. To estimate ***w***_*v*_, L_2_-regularized least-squares estimation was used while the regularization parameter λ was determined based on five-fold cross-validation.

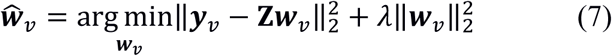

To train this linear regression model, we used the fMRI data acquired during the training movie. The model training was performed separately for the two feature models (the RNN and the CNN) using the same training algorithm. Afterwards, we used the trained encoding models to predict cortical fMRI responses to the independent testing movie. The prediction accuracy was quantified as the temporal correlation (*r*) between the observed and predicted responses at each voxel. As in our previous studies (Wen et al., 2016, 2017a, 2017b), the statistical significance of the prediction accuracy was evaluated voxel by voxel with a block-permutation test (Adolf et al., 2014) corrected at the false discovery rate (FDR) *q* < 0.01.

Given the dimension reduction of the feature space, 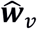 described the contributions to voxel *v* from individual basis vectors in the reduced feature space (i.e. columns of **B** in Eq. (5)). Since the dimension reduction was through linear transform, the voxel-wise encoding models (Eq. (6)) could be readily rewritten with the regressors specific to individual units (instead of basis vectors) in the RNN (or the CNN). In this equivalent encoding model, the regression coefficients, denoted as 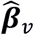, reported the contribution from every unit to each voxel, and could be directly computed from 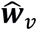 as below.

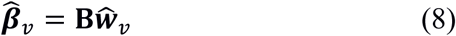

For each voxel, we further identified a subset of units in the RNN that contributed to the voxel’s response relatively more than other units. To do so, the half of the maximum in the absolute values of 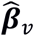 was taken as the threshold. Those units, whose corresponding regression coefficients had absolute values greater than this threshold, were included in a subset (denoted as **I**_*v*_) associated with voxel *v*.

### Model Evaluation and Comparison

After training them using the same training data and the same training algorithms, we compared the encoding models based on the RNN and those based on the CNN. For this purpose, the encoding performance was evaluated as the accuracy of predicting the cortical responses to every session of the testing movie. The prediction accuracy was measured as the temporal correlation (*r*) and then was converted to a z score by Fisher’s z-transformation. For each voxel, the z score was averaged across different movie sessions and different subjects, and the difference in the average z score between the RNN and the CNN was computed voxel by voxel. Such voxel-wise difference was evaluated for statistical significance using the paired t-test across different movie sessions and different subjects (p < 0.01). The differences were also assessed at different ROIs, which were defined based on the cortical parcellation (Glasser et al., 2016), and evaluated for statistical significance using the paired t-test across voxels (p < 0.01).

We also compared the encoding performance against the “noise ceiling”, or the upper limit of the prediction accuracy (Nili et al., 2014). The noise ceiling was lower than 1, due to the fact that the measured fMRI data contained ongoing noise or activity unrelated to the external stimuli, and thus the measured data could not be entirely predictable from the stimuli even if the model were perfect. As described elsewhere (Kay et al., 2013), the response (evoked by the stimuli) and the noise (unrelated to the stimuli) were assumed to be additive and independent and follow normal distributions. Such response and noise distributions were estimated from the data. For each subject, the testing movie was presented ten times. For each voxel, the mean of the noise was assumed to be zero; the variance of the noise was estimated as the mean of the standard errors in the data across the 10 repetitions; the mean of the response was taken as the voxel signal averaged across the 10 repetitions, and the variance of the response was taken as the difference between the variance of the data and the variance of the noise. From the estimated signal and noise distributions, we conducted Monte Carlo simulations to draw samples of the response and the noise, and to simulate noisy data by adding the response and noise samples. The correlation between the simulated response and noisy data was calculated for each of the 1,000 repetitions of simulation, yielding the distribution of noise ceilings at each voxel or ROI.

### Mapping the cortical hierarchy for spatiotemporal processing

We also used the RNN-based encoding models to characterize the functional properties of each voxel, by summarizing the fully-observable properties of the RNN units that were most predictive of that voxel. As aforementioned, each voxel was associated with a subset of RNN units **I**_*v*_. In this subset, we calculated the percentage of the units belonging to each of the four layers (indexed by 1 through 4) in the RNN, multiplied the layer-wise percentage by the corresponding layer index, and summed the result across all layers to yield a number (between 1 and 4). This number was assigned to the given voxel *v*, indicating this voxel’s putative “level” in the visual hierarchy. Mapping the voxel-wise level revealed the hierarchical cortical organization for spatiotemporal visual processing.

### Estimating Temporal Receptive Windows

We also quantified the temporal receptive window (TRW) at each voxel *v* by summarizing the “temporal dependency” of its contributing units **I**_*v*_ in the RNN. For each unit *i* ∈ **I**_*v*_, its forget gate, denoted as 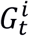, controlled the memory storage vs. retrieval at each moment *t*. For simplicity, let us define a “remember” gate, 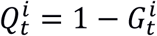, to act oppositely as the forget gate. From Eq. (1), the current state (or unit activity) 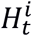 was expressed as a function of the past input {**x**_*t*-τ_|1 ≤ τ ≤ *t*}.

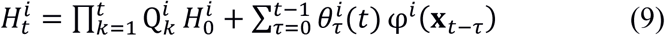

where 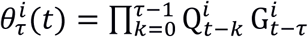. In Eq. (9), the first term was zero given the initial state 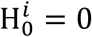 The second term was the result of applying a time-variant filter 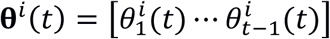 to the time series of the spatial representation {φ ^*i*^(**x**_*t*_)}_∀*t*_ extracted by the CNN from every input frame {**x**_*t*_}_∀*t*_. In this filter, each element 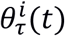 reflected the effect of the past visual input **x**_*t*-τ_ (with an offset τ) on the current state 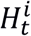. As it varied in time, we averaged the filter **θ**^*i*^(*t*) across time, yielding 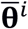 to represent the average temporal dependency of each unit *i*.

From the observable temporal dependency of every unit, we derived the temporal dependency of each voxel by using the linear unit-to-voxel relationships established in the encoding model. For each voxel *v*, the average temporal dependency was expressed as a filter 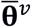, which was derived as the weighted average of the filters associated with its contributing RNN units 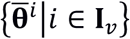, as in Eq. (10).

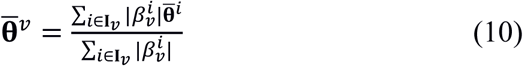

Of 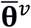, the elements 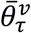 delineated the dependency of the current response at voxel *v* on the past visual input with an offset τ prior to the current time. The accumulation of temporal information was measured as the sum of 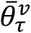 across different offsets in a given time window. The window size that accounted for 95% of the accumulative effect integrated over an infinite past period was taken as the TRW for voxel *v*. In the level of ROIs, the TRW was averaged across voxels within each pre-defined ROI. The difference in TRW between different ROIs was evaluated using two-sample t-tests (p < 0.01).

### Spectral Analysis of Forget Gate Dynamics

We also characterized the temporal fluctuation of the forget gate at each unit in the RNN. As the forget gate behaved as a switch for controlling, moment by moment, how information was stored into vs. retrieved from process memory. As such, the forget-gate fluctuation reflected the dynamics of process memory in the RNN given natural video inputs.

To characterize the forget-gate dynamics, its power spectral density (PSD) was evaluated. The PSD followed a power-law distribution that was fitted with a descending line in the double-logarithmic scale. The slope of this line, or the power-law exponent (PLE) (Miller et al., 2009; Wen and Liu, 2016), characterized the balance between slow (low-frequency) and fast (high-frequency) dynamics. A higher PLE implied that slow dynamics dominated fast dynamics; a lower PLE implied the opposite. After the PLE was evaluated for each unit, we derived the PLE for each voxel *v* as a weighted average of the PLE of every unit *i* that contributed to this voxel (*i* ∈ **I**_*v*_), in a similar way as expressed in Eq. (10).

## RESULTS

### RNN learned video representations for action recognition

We used a recurrent neural network (RNN) to model and predict cortical fMRI responses to natural movie stimuli. This model extended a pre-trained CNN (VGG16) (Simonyan and Zisserman, 2014a) by adding recurrent connections to different layers in the CNN (**Fig. 1**). While fixing the CNN, the weights of the recurrent connections were optimized by supervised learning with >13,000 labeled videos from 101 action categories (Soomro et al., 2012). After training, the RNN was able to categorize independent test videos with a 76.7% top-1 accuracy. This accuracy was much higher than the 65.09% accuracy obtained with the CNN, and close to the 78.3% accuracy obtained with the benchmark RNN model (Ballas et al., 2015).

Unlike the CNN, the RNN explicitly embodied a network architecture to learn hierarchically organized video representations for action recognition. When taking isolated images as the input, the RNN behaved as a feedforward CNN for image categorization. In other words, the addition of recurrent connections enabled the RNN to recognize actions in videos, without losing the already learned ability for recognizing objects in images.

### RNN better predicted cortical responses to natural movies

Accompanying its enriched AI, the RNN learned to utilize the temporal relationships between video frames, whereas the CNN treated individual frames independently. We asked whether the RNN constituted a better model of the visual cortex than the CNN, by evaluating and comparing how well these two models could predict cortical fMRI responses to natural movie stimuli. The prediction was based on voxel-wise linear regression models, through which the representations of the movie stimuli, as extracted by either the RNN or the CNN, were projected onto each voxel’s response to the stimuli. Such regression models were trained and tested with different sets of video stimuli (12.4 hours or 2.4 hours for training, 40 minutes for testing) to ensure unbiased model evaluation and comparison. Both the RNN and the CNN explained significant variance of the movie-evoked response for widespread cortical areas (**Fig. 2A & 2B**). The RNN consistently performed better than the CNN, showing higher prediction accuracy for nearly all visual areas (**Fig. 2D**), especially for cortical locations along the dorsal visual stream (**Fig. 2C**). The response predictability given the RNN was about half of the “noise ceiling” – the upper limit by which the measured response was predictable given the presence of any ongoing “noise” or activity unrelated to the stimuli (**Fig. 2D**). This finding was consistently observed for each of the three subjects (**Fig. 3**).

**Figure 2.**
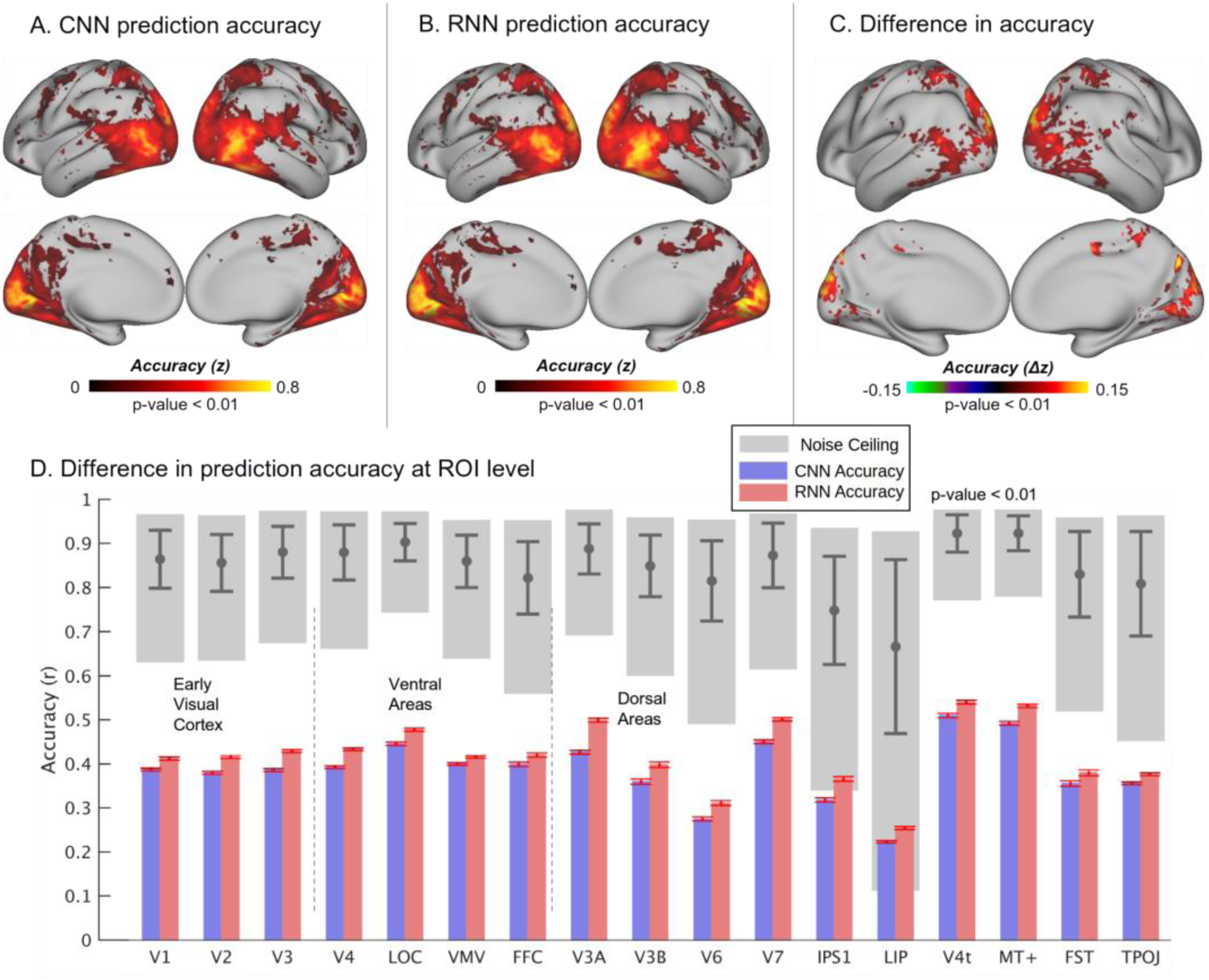
Prediction accuracies of the cortical responses to novel movie stimuli. **A)** Performance of the CNN-based encoding model, averaged across testing movie sessions and subjects. **B**) Performance of the RNN-based encoding model, averaged across testing movie sessions and subjects. **C**) Significant difference in the performance between the RNN and CNN. The values of difference were computed as subtracting **A**) from **B**). **D**) Comparison of performances at different ROIs with noise ceilings. The accuracy at each ROI is the voxel mean within the region, where the red bars indicate the standard error of accuracies across voxels. The gray blocks indicate the lower and upper bounds of the noise ceilings, and the gray bars indicate the mean and standard deviation of the noise ceilings at each ROI.

**Figure 3.**
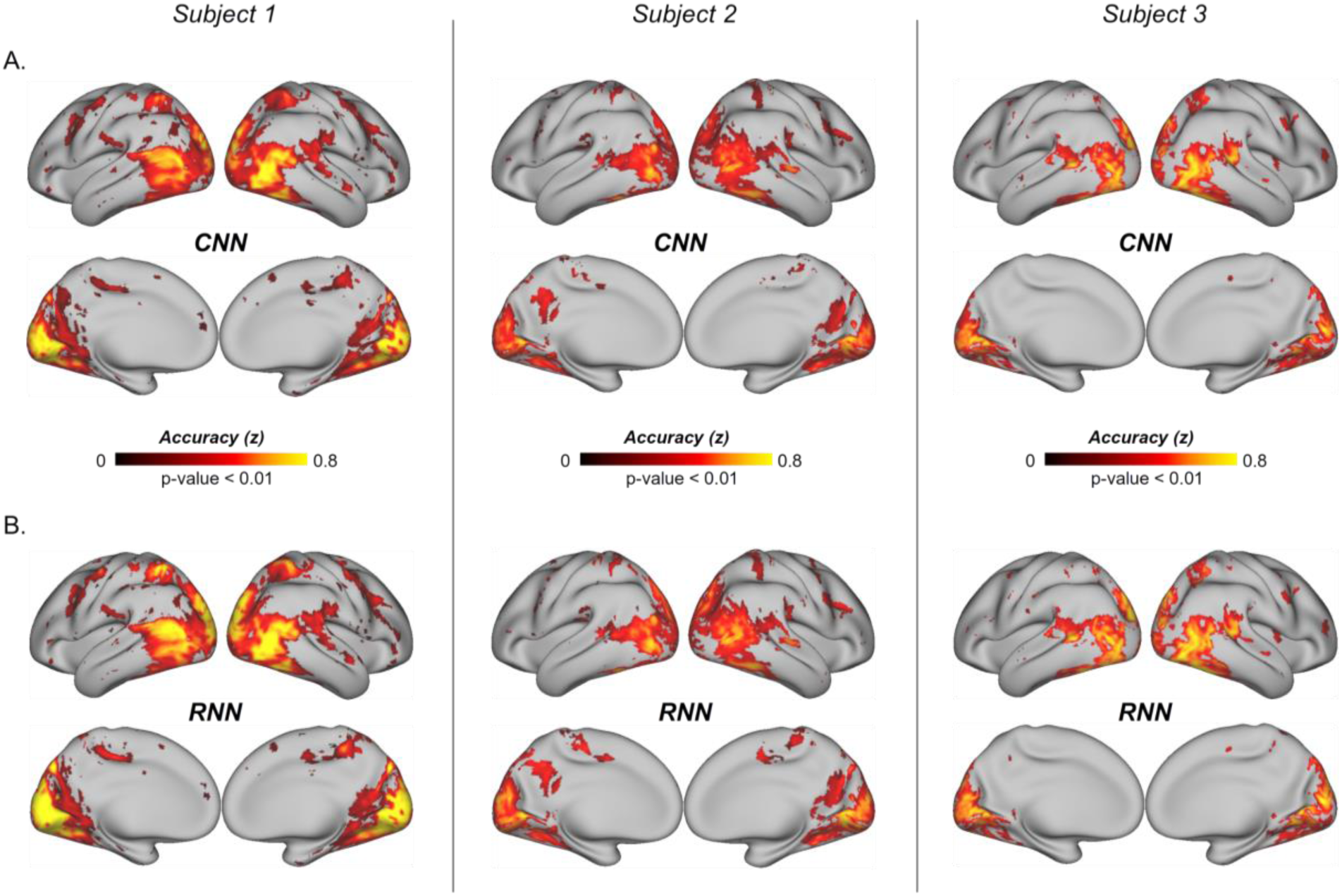
Prediction accuracies of the cortical responses to novel movie stimuli for individual subjects. **A)** Performance of the CNN-based encoding model, averaged across testing movie sessions. **B**) Performance of the RNN-based encoding model, averaged across testing movie sessions.

### RNN revealed a gradient in temporal receptive windows (TRWs)

Prior studies have shown empirical evidence that visual areas were hierarchically organized to integrate information not only in space (Kay et al., 2013), but also in time (Hasson et al., 2008). Units in the RNN learned to integrate information over time through the unit-specific “forget gate”, which controlled how past information shaped processing at the present time. Through the linear model that related RNN units to each voxel, the RNN’s temporal “gating” behaviors were passed from units to voxels in the brain. As such, this model allowed to characterize the TRWs, in which past information was carried over and integrated over time to affect and explain the current response at each specific voxel or region.

**Fig. 4A** shows the response at each given location as the accumulative effect integrated over a varying period (or window) prior to the current moment. On average, the response at V1 reflected the integrated effects over the shortest period, suggesting the shortest TRW at V1. Cortical areas running down the ventral or dorsal stream integrated information over progressively longer TRWs (**Fig. 4A**). Mapping the voxel-wise TRW showed a spatial gradient aligned along the visual streams, suggesting a hierarchy of temporal processing in the visual cortex (**Fig. 4B**). In the ROI level, the TRWs were significantly shorter for early visual areas than those for higher-order ventral or dorsal areas; and the dorsal areas tended to have longer TRWs than the ventral areas (**Fig. 4C**). While Fig. 4 shows the results for Subject 1, similar results were also observed in the other subjects (**Fig. S1** and **Fig. S2**). We interpret the TRW as a measure of the average capacity of process memory at each cortical location involved in visual processing.

**Figure 4.**
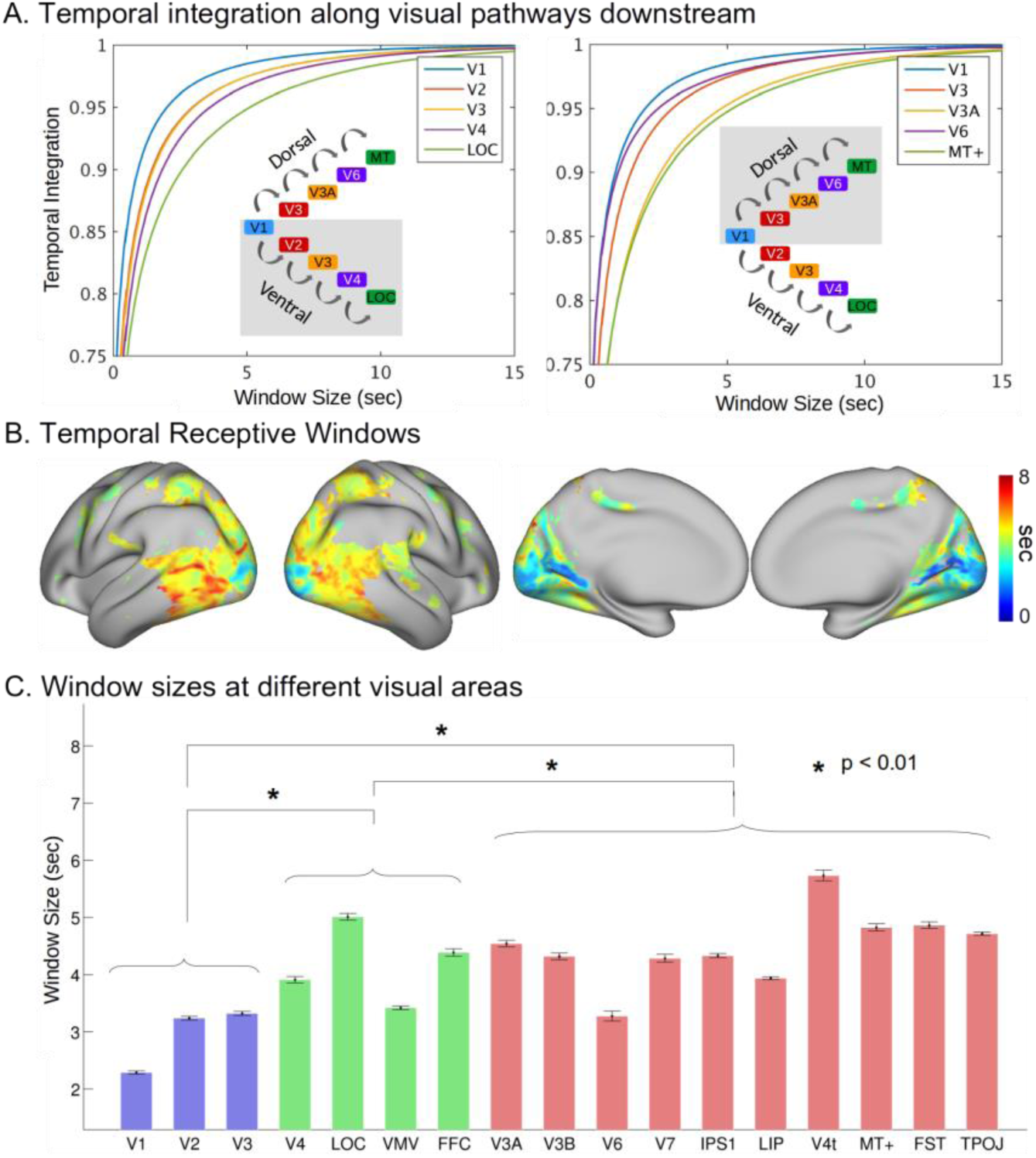
Model-estimated TRWs in the visual cortex of Subject 1. **A)** The accumulation of information at different ROIs along ventral and dorsal streams. Window size represents the period to the past, and temporal integration indicates the relative amount of accumulated information. **B**) The cortical map of TRWs estimated by the RNN. The color bar indicates the window sizes at individual voxels. **C**) Average TRWs at individual ROIs. The blue bars represent the early visual cortex, the green bars the ventral areas, and the red bars the dorsal areas. The black error bars indicate the standard errors across voxels.

### RNN revealed the slow vs. fast dynamics of process memory

In the RNN, the forget gate varied from moment to moment, indicating how the past vs. current information was mixed together to determine the representation at each moment. Given the testing movie stimuli, the dynamics of the forget gate was scale free, showing a power-law relationship in the frequency domain. The power-law exponent (PLE) reported on the balance between slow and fast dynamics: a higher exponent indicated a tendency for slow dynamics, and a lower exponent indicated a tendency for fast dynamics.

After projecting the PLEs from units to voxels, we mapped the distribution of the voxel-wise PLE to characterize the dynamics of process memory (Hasson et al., 2015) at each cortical location. As shown in **Fig. 5**, the PLE was lower in early visual areas, but became increasingly larger along the downstream pathways in higher-order visual areas. Such trend was analogous to the gradient in TRWs (**Fig. 4B**), where the TRWs were shorter in early visual areas and longer in higher-order visual areas. In general, lower PLEs were associated with areas with shorter TRWs; higher PLEs were associated with areas with longer TRWs.

**Figure 5.**
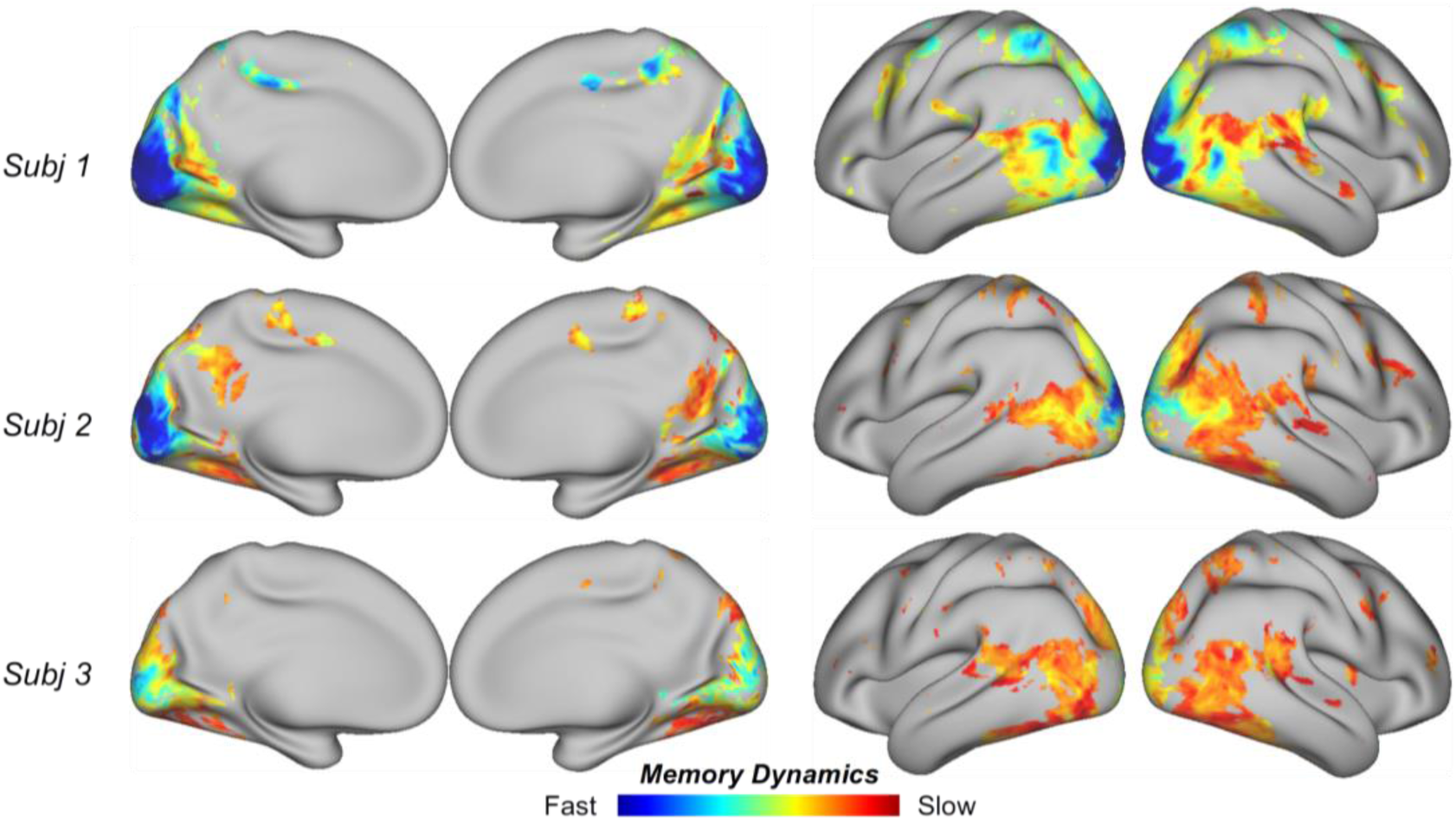
Model-estimated memory dynamics in the visual cortex. Consistent across subjects, lower PLEs are associated early visual areas, and higher PLEs are associated with later stages of visual processing.

### RNN revealed the cortical hierarchy of spatiotemporal processing

CNNs revealed the hierarchical organization of spatial processing in the visual cortex (Güçlü and van Gerven, 2015a; Wen et al., 2016; Eickenberg et al., 2017; Horikawa and Kamitani, 2017). By using the RNN as a network model for spatiotemporal processing, we further mapped the hierarchical cortical organization of spatiotemporal processing. To do so, every voxel, where the response was predictable by the RNN, was assigned with an index, ranging continuously between 1 and 4. This index reported the “level” that a voxel was involved in the visual hierarchy: a lower index implied an earlier stage of processing; a higher index implied a later stage of processing. The topography of the voxel-wise level index showed a cortical hierarchy (**Fig. 6**). Locations from striate to extra-striate areas were progressively involved in early to late stages of processing the information in both space and time.

**Figure 6.**
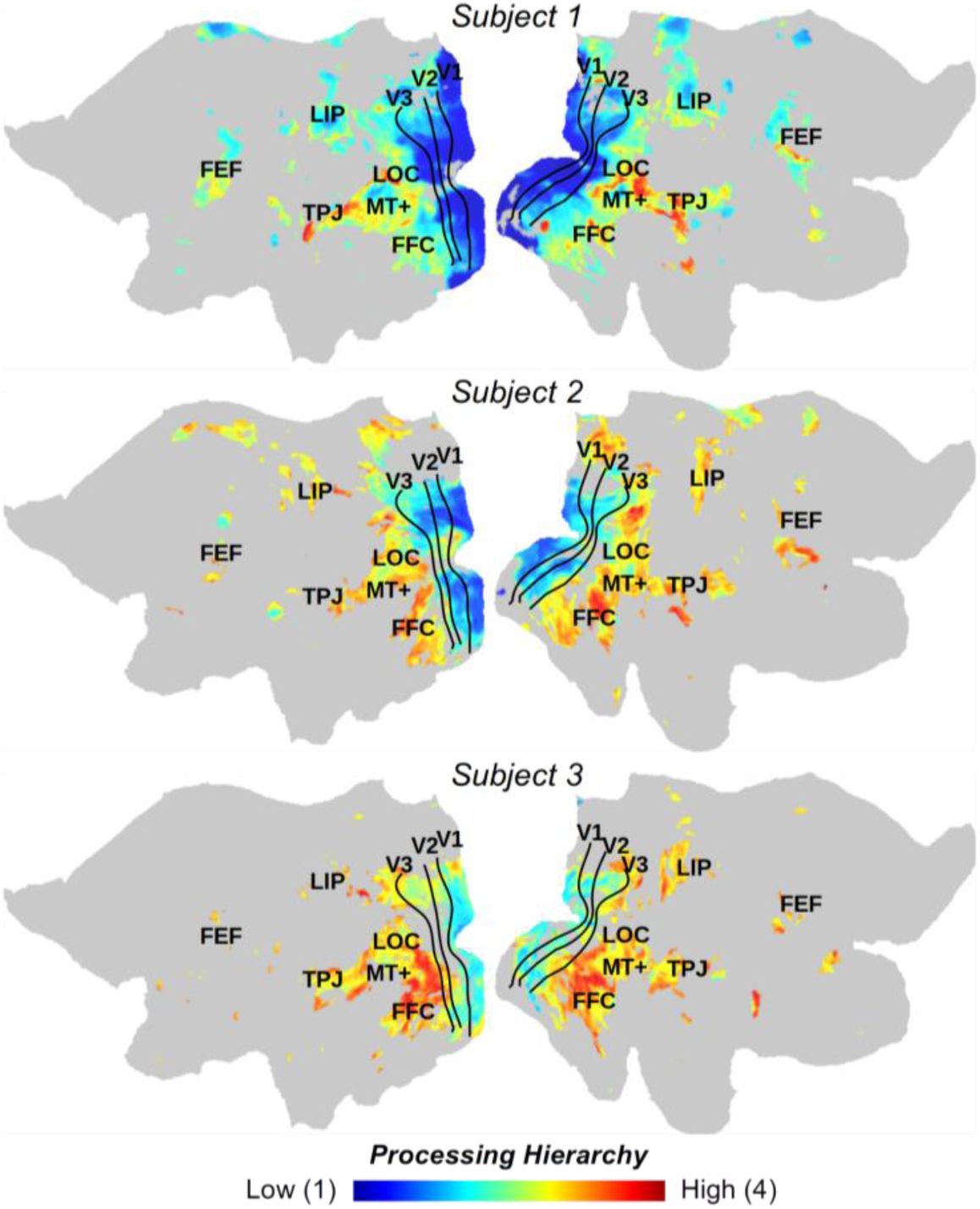
Model-estimated hierarchical organization of spatiotemporal processing. Consistent across subjects, lower layer indices are assigned to early visual areas, and higher layer indices are assigned to later stages of visual processing. The color bar indicates the range of layer assignment, from layer 1 to 4.

## DISCUSSION

Here, we designed and trained a recurrent neural net (RNN) to learn video representations for action recognition, and to predict cortical responses to natural movies. This RNN extended from a pretrained CNN by adding layer-wise recurrent connections to allow visual information to be remembered and accumulated over time. In line with the hypothesis of process memory (Hasson et al., 2015), such recurrent connections formed a hierarchical and distributed model of memory as an integral part of the network for processing dynamic and natural visual input. Compared to the CNN, the RNN supported both image and action recognition, and better predicted cortical responses to natural movie stimuli at all visual areas, especially those along the dorsal stream. More importantly, the RNN provided a fully-observable computational model to characterize and map temporal receptive windows, dynamics of process memory, and a cortical representational hierarchy for dynamic natural vision.

### A network model of process memory

Our work was in part inspired by the notion of “process memory” (Hasson et al., 2015). In this notion, memory is a continuous and distributed process as an integral part of information processing, as opposed to an encapsulated functional module separate from the neural circuits that process sensory information. Process memory provides a mechanism for the cortex to process the temporal information in natural stimuli, in a similarly hierarchical way as cortical processing of spatial information (Hasson et al., 2015). As explored in this study, the RNN uses an explicit model of process memory to account for dynamic interactions between incoming stimuli and the internal states of the neural network, or the state-dependent computation (Buonomano and Maass, 2009). In the RNN, the “forget gate” controls, separately for each unit in the network, how much its next state depends on the incoming stimuli vs. its current state. As such, the forget gate behaves as a switch of process memory to control how much new information should be stored into memory and how much history information should be retrieved from memory. This switch varies moment to moment, allowing memory storage and retrieval to occur simultaneously and continuously.

As demonstrated in this study, this model of process memory could be trained, with supervised learning, for the RNN to classify videos into action categories with a much higher accuracy than the CNN without any mechanism for temporal processing. It suggests that integrating process memory to a network of spatial processing indeed makes the network to be capable of spatiotemporal processing, as implied in previous theoretical work (Buonomano and Maass, 2009).

### From theoretical modeling to empirical evidence of process memory

A unique contribution of this study is that computational modeling of process memory is able to explain previous empirical evidence for process memory. One of the strongest evidence for process memory is that the cortex organizes a topography of temporal receptive window (Hasson et al., 2008; Honey et al., 2012), which may be interpreted as the voxel-wise capacity of process memory. To probe the TRW, an experimental approach is to scramble the temporal structure of natural stimuli in multiple timescales and measure their resulting effects on cortical responses (Hasson et al., 2008). The TRW measured in this way increases orderly from early sensory areas to higher-order perceptual or cognitive areas (Hasson et al., 2015), suggesting a hierarchical organization of temporal processing. With this approach, the brain is viewed as a “black box” and is studied by examining its output given controlled perturbations to its input.

In this study, we have reproduced the hierarchically organized TRW by using a model-driven approach. The RNN tries to model the inner-working of the visual cortex as a computable system, such that the system’s output can be computed from its input. If the model uses the same computational and organizational principles as does the brain itself, the model’s output should match the brain’s response given the same input (Wu et al., 2006; Naselaris et al., 2011). By “matching”, we do not mean that the unit activity in the model should match the voxel response in the brain with one-to-one correspondence, but up to linear transform (Yamins and Di Carlo, 2016), because it is unrealistic to exactly model the brain. This approach allows to test computational models against experimental findings. The fact that the model of process memory explains the topography of TRW (i.e. the hallmark evidence for process memory), lends synergistic support to process memory as a fundamental principle for spatiotemporal processing of natural visual stimuli.

### RNN extends CNN as both a brain model and an AI

Several recent studies explored deep-learning models as predictive models of cortical responses during natural vision (Yamins et al., 2014; Khaligh-Razavi et al., 2014; Güçlü and van Gerven, 2015a, b; Wen et al., 2016, 2017a, 2017b; Cichy et al., 2016; Eickenberg, et al., 2017; Horikawa and Kamitani, 2017). Most of the prior studies used CNNs that extracted spatial features to support image recognition, and demonstrated the CNN as a good model for the feedforward process along the ventral visual stream (Yamins et al., 2014; Khaligh-Razavi et al., 2014; Güçlü and van Gerven, 2015a; Eickenberg, et al., 2017). In our recent study (Wen et al., 2016), the CNN was further found to be able to partially explain the dorsal-stream activity in humans watching natural movies; however, the predictive power of the CNN was lesser in the dorsal stream than in the ventral stream. Indeed, the dorsal stream is known for its functional roles in temporal processing and action recognition in vision (Goodale and Milner, 1992; Rizzolatti and Matelli, 2003; Shmuelof and Zohary, 2005). It is thus expected that the limited ability of the CNN for explaining the dorsal-stream activity is due to its lack of any mechanism for temporal processing.

Extending from the CNN, the RNN established in this study offered a network mechanism for temporal processing, and improved the performance in action recognition. Along with this enhanced performance towards humans’ perceptual ability, the RNN also better explained human brain activity than did the CNN (**Fig. 2**). The improvement was more apparent in areas along the dorsal stream than those along the ventral stream (**Fig. 2**). It is worth noting that when the input is an image rather than a video, the RNN behaves as the CNN to support image classification. In other words, the RNN extends the CNN to learn a new ability (i.e. action recognition) without losing the already learned ability (i.e. image recognition). On the other hand, the RNN, as a model of the visual cortex, improves its ability in predicting brain activity not only at areas where the CNN falls short (i.e. dorsal-stream), but also at areas where the CNN excels (i.e. ventral-stream). As shown in this study, the RNN better explained the dorsal stream, without losing the already established ability to explain the ventral stream (**Fig. 2**). This brings us to a perspective about developing brain models or brain-inspired AI systems. As humans continuously learn from experiences to support different intelligent behaviors, it is desirable for an AI model to continuously learn to expand capabilities while keeping existing capabilities. When it is also taken as a model of the brain, this AI model should be increasingly more predictive of brain responses at new areas, while remaining its predictive power at areas where the model already predicts well. This perspective is arguably valuable for designing a brain-inspired system for continuous learning as does the brain itself.

### Comparison with related prior work

Other than RNN, a 3-dimensional (3-D) CNN may also learn spatiotemporal features for action recognition of videos (Tran et al., 2014). A 3-D CNN shares the same computational principle as an otherwise 2-D CNN, except that the input to the former is a time series of video frames with a specific duration, whereas the input to the latter is a single video frame or image. Previously, the 3-D CNN was shown to explain cortical fMRI responses to natural movie stimuli (Güçlü and van Gerven, 2015b). However, it is unlikely that the brain works in a similar way as a 3-D CNN. The brain processes visual information continuously delivered from 2-D retinal input, rather than processing time blocks of 3-D visual input as required for 3-D CNN. Although it is a valid AI model, 3-D CNN is not appropriate for modeling or understanding the brain’s mechanism of dynamic natural vision.

It is worth noting the fundamental difference between the RNN model in this study and that in a recently published study (Güçl ü and van Gerven, 2017). Here, we used the RNN as a feature model or the model of the visual cortex, whereas Güçlü and van Gerven used the RNN as the response model in an attempt to better describe the complex relationships between the CNN and the brain. Although a complex response model is potentially useful, it defeats our purpose of seeking a computational model that matches the visual cortex up to linear transform. It is, has been, and will be, our intention to find a model that shares similar computing and organization principles as the brain. Towards this goal, the response model needs to as simple as possible, independent of the visual input, and with canonical or independently defined HRF.

### Future Directions

The focus of this study is on vision. However, the RNN is expected to be useful, or even more useful, for modeling other perceptual or cognitive systems beyond vision. RNNs have been successful in computer vision (Donahue et al., 2015), natural language processing (Hinton et al., 2012; Mikolov et al., 2010), attention (Mnih et al., 2014; Xu et al., 2015; Sharma et al., 2015), memory (Graves et al., 2014), and planning (Zaremba and Sutskever, 2015). It is conceivable that such RNNs would set a good starting point to model the corresponding neural systems, to facilitate the understanding of the network basis of complex perceptual or cognitive functions.

The RNN offers a computational account of temporal processing. If the brain performs similar computation, how is it implemented? The biological implementation of recurrent processing may be based on lateral or feedback connections (Lamme et al., 1998; Kafaligonul et al., 2015). The latter is of particular interest, since the brain has abundant feedback connections to exert top-down control of feedforward processes (Itti et al., 1998; de Fockert et al., 2001). However, the feedback connections are not taken into account in this study, but may be incorporated into the models in the future by using such brain principles as predictive coding (Rao and Ballard, 1999) or the free-energy principle (Friston, 2010). Recent efforts along this line are promising (Lotter et al., 2016; Canziani and Culurciello, 2017) to merit further investigation.

## Acknowledgements

This work was supported in part by NIH R01MH104402. The authors would like to recognize the inputs from Dr. Eugenio Culurciello on the discussions of deep neural networks.

